# Antisense oligonucleotide-based drug development for Cystic Fibrosis patients carrying the 3849+10kb C-to-T splicing mutation

**DOI:** 10.1101/2021.02.14.431123

**Authors:** Yifat S. Oren, Michal Irony-Tur Sinai, Anita Golec, Ofra Barchad-Avitzur, Venkateshwar Mutyam, Yao Li, Jeong Hong, Efrat Ozeri-Galai, Aurélie Hatton, Joel Reiter, Eric J. Sorscher, Steve D. Wilton, Eitan Kerem, Steven M. Rowe, Isabelle Sermet-Gaudelus, Batsheva Kerem

**Author notes:** These two authors contributed equally (Yifat S. Oren, Michal Irony-Tur Sinai). These two authors contributed equally (Anita Golec, Ofra Barchad-Avitzur). **Address correspondance to**: Batsheva Kerem, Department of Genetics, The Life Sciences Institute, The Hebrew University, Edmond J Safra Campus, Givat Ram, Jerusalem, Israel 91904.

## Abstract

Antisense oligonucleotide (ASO)-based drugs for splicing modulation were recently been approved for various genetic diseases with unmet need. Here we aimed to develop an ASO-based splicing modulation therapy for Cystic Fibrosis (CF) patients carrying the 3849+10kb C-to-T splicing mutation in the CFTR gene. We have screened, in FRT cells expressing this mutation, ~30 ASOs chemically modified with 2′-O-Methyl on a phosphrothioate backbone, targeted to prevent the recognition and inclusion of a cryptic exon generated due to the mutation. The screening identified five ASO candidates able to promote CFTR correct splicing and rescue channel activity. Further analyses in well differentiated primary human nasal and bronchial epithelial cells (HNEs, HBEs), derived from patients carrying at least one 3849+10kb C-to-T allele, led to the identification of a highly potent lead ASO. The ASO was efficiently delivered by free uptake into patients’ HNEs and HBEs and completely restored CFTR function to wild type levels in cells from a homozygous patient and led to 43±8% of wild type levels in cells from various heterozygous patients. Optimized efficiency was further obtained with 2’-Methoxy Ethyl chemical modification. The results demonstrate the therapeutic potential and clinical benefit of ASO-based splicing modulation for genetic diseases caused by splicing mutations.

## Introduction

Cystic fibrosis (CF) is a life-shortening multi-organ genetic disease affecting ~ 90,000 individuals worldwide, caused by mutations in the cystic fibrosis transmembrane conductance regulator (CFTR) gene that encodes a chloride (Cl^−^) channel located at the surface of epithelial cells resulting in impaired ion transport across tissues of the exocrine system [reviewed in (1)]. The impaired CFTR activity alters electrolytes and hydration balance across epithelia, causing the accumulation of thick mucus in the lung bronchial tree leading to a chronic progressive lung disease, which is the major cause of morbidity and mortality [reviewed in (1)].

The last decade has witnessed developments of genotype-specific, targeted drugs that improve CFTR protein folding, stability and gating defects, leading to increased amounts of mutant CFTR reaching the cell surface and to restoration of the ion transport. These CFTR modulators offer therapeutic opportunities mainly for patients that carry gating mutations or the F508del mutation [reviewed in (2)]. Still, there are CF patients carrying CFTR mutations for which the current CFTR modulators are unlikely to provide a clinical benefit. Among these are patients carrying mutations abrogating the production of CFTR proteins, such as deletions, nonsense and splicing mutations. Among the ~2000 reported CFTR sequence variations, a significant fraction (10-15%) affect splicing of the precursor messenger RNA (pre-mRNA), by either creating or abolishing canonical splice sites, commonly leading to skipping over the exon. There is another group of mutations altering exonic and intronic regulatory splicing motifs throughout the gene (3), leading to variable levels of both aberrantly and correctly spliced transcripts from these mutated alleles. This group includes the splicing mutations 3849+10kb C-to-T (c.3717+12191C-to-T), 1811+1.6 kb A-to-G (c.1679+1634A>G), 3272-26A-to-G (c.3140-26A>G) and IVS8-5T, 2789+5G-to-A (c.2657+5G>A) [reviewed in (4)]. Most patients carrying splicing mutations are pancreatic sufficient, however, clinical studies show that their lung function is variable and is similar to that observed among patients with CF carrying minimal function mutations (https://cftr2.org/)(5–7).

The 3849+10kb C-to-T splicing mutation, generates an aberrant 5′ splice site, deep in intron 22 of the CFTR pre-mRNA. This activates a cryptic 3′ splice site 84 nucleotides upstream, resulting in the inclusion of 84 intronic nucleotides that constitute a cryptic exon in the CFTR mRNA (8). This 84bp cryptic exon contains an in-frame stop codon, leading to degradation of a significant fraction of the mRNA by the nonsense-mediated mRNA decay (NMD) mechanism, as well as to the production of truncated non-functional CFTR proteins (9). Importantly, the 3849+10 kb C-to-T mutation does not alter the wild-type (WT) splice sites and can enable the generation of both aberrantly and correctly spliced transcripts. Since the normal CFTR splice site sequences are intact, the involved pre-mRNA retains the potential for normal splicing, if usage of the aberrant splice sites could be inhibited. This mutation is the 7^th^ most common CFTR mutation in the US and 8^th^ in Europe, carried by >1400 CF patients worldwide (10, 11). In several populations, the mutation is highly prevalent, such as in Ashkenazi Jews and CF patients in Slovenia, Poland and Italy. Since this mutation is associated with reduced amount of normal CFTR, clinical trials investigated the effect of the modulator ivacaftor alone or together with tezacaftor and showed a modest clinical benefit (12, 13). Therefore, another approach is required in order to restore the CFTR function and significantly improve the disease in patients carrying alternative splicing mutations.

Importantly, a correlation between lung disease severity as measured by lung function and the level of correctly spliced CFTR transcripts was found for patients carrying various splicing mutations, including the 3849+10kb C-to-T mutation [(8, 14–17) as reviewed in (18)]. This correlation is found also among patients with the same genotype (16, 17). The ability of the splicing machinary to act as a disease modifier was demonstrated in several models of genetic diseases caused by splicing mutations [reviewed in (19, 20)]. For example, overexpression of splicing factors was able to increase the level of correctly spliced CFTR RNA transcribed from the 3849+10 kb C-to-T allele and to promote the restoration of CFTR channel function (21). These observations, which highlight the potential of splicing modulation as a therapeutic approach, and the therapeutic need which still exists for CF patients carrying splicing mutations (12, 13), encouraged us to develop drug candidates with a specific splice-switching potential.

A specific therapeutic approach for splicing modulation is based on the administration of single-stranded short Antisense Oligonucleotides (ASOs) designed to hybridize to specific elements within target RNAs [reviewed in (22, 23)]. Splice switching ASO-based therapies are designed to inhibit or activate specific splicing events by a steric blockade of the recognition of specific splicing elements and preventing the recruitment of effectors to these sites. The potential of ASOs to modulate the splicing pattern generated due to CFTR splicing mutations was shown in cellular systems overexpressing full-length mutated CFTR cDNA constructs. ASO transfection of epithelial cell cultures, expressing a CFTR cDNA vector harbouring a mini-intron 22 with the 3849+10 kb C-to-T locus, enhanced normal CFTR splicing and increased the production of normally processed CFTR proteins (24). Similarly, ASO transfection of epithelial cells expressing a cDNA harbouring the c.2657+5G>A (2789+ 5G>A) splicing mutation, which causes the generation of CFTR transcripts lacking exon 16, increased the amount of correctly spliced CFTR proteins localized at the plasma membrane and restored CFTR function (25). The ability of ASOs to modulate the splicing of the endogenous 3849+10kb C-to-T allele was recently demonstrated by Michaels et al., showing that transfecting primary Human Bronchial Epithelial cells (HBEs) with a phosphorodiamidate morpholino oligomer (PMO) targeted to mask the cryptic splice site was able to block aberrant splicing and to improve CFTR function (26).

ASO-based drugs modulating splicing are already approved for Spinal muscular atrophy (SMA) and Duchenne muscular dystrophy (DMD) [reviewed in (27, 28)]. The exciting clinical data suggests that ASO-mediated splicing modulation is able to improve protein function and slow disease progression. In light of this data, modulating the level of correctly spliced CFTR transcripts using an ASO-based approach has a great therapeutic potential for CF patients.

Here we focused on the development of drug candidates for patients carrying the 3849+10Kb C-to-T splicing mutation, using chemically modified ASOs targeted to prevent the recognition of splicing elements involved in the cryptic exon inclusion. We have identified a lead ASO able to significantly increase the level of correctly spliced mRNA and restore the production of normal and functional CFTR channels by a free ASO uptake in well differentiated polarized Human Nasal Epithelial (HNE) and HBE cells, from patients carrying the 3849+10Kb C-to-T allele. Our promising results are aimed to serve as a basis for clinical evaluation of the lead ASO.

## Results

### ASO treatment promotes CFTR correct splicing in HEK293 cells over-expressing the 3849+10kb C-to-T mutation

In the present study, we aimed to develop an ASO-based therapy that will prevent the recognition of the 84bp cryptic exon and promote correct splicing. First, we designed two ASOs [ASO84-1 and ASO84-4, 25 nucleotides (nt) in length] targeting the junctions between the 84bp cryptic exon and flanking intronic sequences (Figure 1A). Masking these junctions was aimed to prevent the recognition of the cryptic exon, redirecting the splicing machinery to the authentic splice motifs. For a control ASO we used an oligonucleotide sequence that has no target in the human genome (29). The ASOs were synthesized with the 2′-O-Methyl (2’-OMe) sugar modification, which provides increased affinity to the targeted mRNA, resistance to nucleases and do not elicit cleavage of the hybridized mRNA by RNase H (30, 31). The ASOs had a full phosphorothioate backbone (PS) which enhances uptake through the cell membrane and provides favorable pharmacokinetic properties (31).

**Figure 1.**
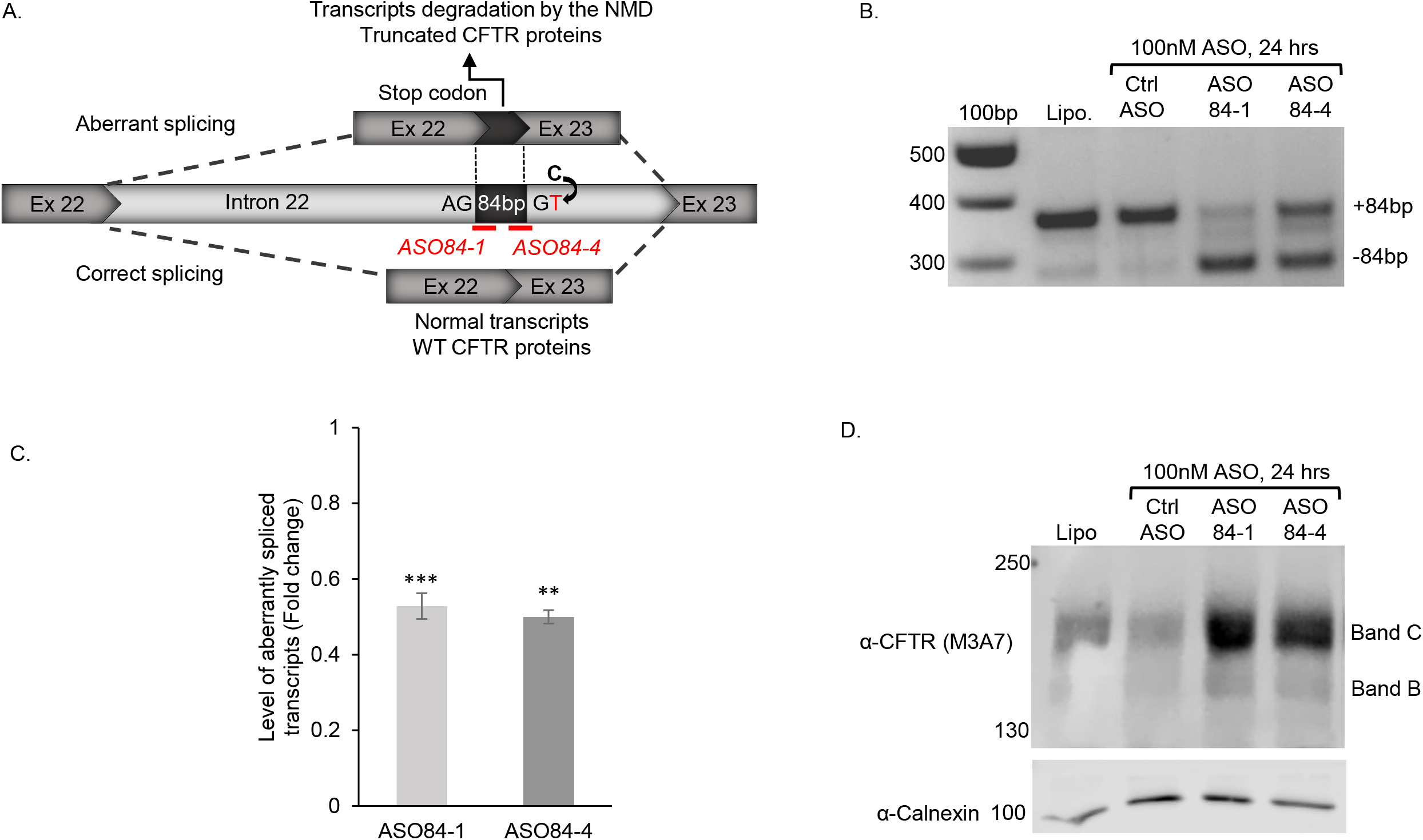
ASO modulation of the aberrant splicing generated from the 3849+10kb C-to-T mutation in HEK293 cells. **A.** A scheme of the splicing pattern generated from the 3849+10kb C-to-T splicing mutation and its effect on the inclusion of the 84 bp cryptic exon in the mature mRNA. The cryptic exon contains an in-frame stop codon, which leads to the production of reduced levels of CFTR transcripts, due to degradation by the NMD mechanism, and truncated nonfunctional CFTR protein. Schematic representation of ASO84-1 and ASO84-4 targeted to the junction sequences between the 84 bp cryptic exon and its flanking intronic sequences is shown (marked in red). **B.** HEK293-3849-mut cells were re-transfected with 100nM of ASO84-1 or ASO84-4 or control ASO for 24 hours. RNA was extracted 24 hours following the second transfection and RT-PCR using primers located in exons 22 and 23 was performed to amplify the aberrantly spliced CFTR transcripts (+84 bp) and the correctly spliced transcripts (−84bp). A representative RT-PCR example, showing both the aberrantly and correctly spliced CFTR transcripts is presented (n=4). Lipofectamine 2000 was used for the mock transfection. **C.** Quantitative analysis of the level of aberrantly spliced CFTR transcripts as measured by RT-qPCR. The values shown are the average fold change (Mean±SEM) from 4 independent experiments relative to cells treated with a control ASO. Values were normalized against transcripts of GUSb gene. Statistical analysis was performed using paired t test (one-tail). **p<0.01, ***p<0.001. **D.** Protein extracts were prepared from the treated HEK293-3849-mut and analyzed by immunoblotting with anti CFTR (M3A7) and anti‐calnexin antibodies. A representative blot is presented (n=4).

In order to study the effect of ASO84-1 and ASO84-4 on the cryptic exon recognition and splicing, we constructed plasmids expressing the full length CFTR cDNA into which sequences from intron 22 containing the 84 bp cryptic exon were introduced. The 3849+10kb C-to-T mutation or the WT sequence and relevant flanking intronic sequences, required and sufficient for splicing, were inserted into the plasmids (CFTR-3849-mut and CFTR-3849-WT plasmids, respectively) (Supplementary Figure 1, see Materials and Methods for details). We first used HEK293 cells, which lack endogenous expression of CFTR. The cells were transiently transfected with the CFTR-3849-mut plasmid (HEK293-3849-mut), and 24 hours later were transfected with the ASO. After 24 hours, RNA was extracted and the CFTR splicing pattern was analyzed. The analysis revealed a significant shift in the ratio between the aberrantly and correctly spliced CFTR transcripts (Figure 1B). While HEK293-3849-mut cells, following mock or control ASO transfection, have mainly aberrantly spliced transcripts, transfection with ASO84-1 or ASO84-4 significantly reduced the relative level of aberrant CFTR transcripts (Figure 1B). Quantitative analysis by RT-qPCR showed that both ASO84-1 and ASO84-4 modified the CFTR splicing pattern, leading to a ~50% reduction in the level of aberrantly spliced transcripts (Figure 1C). In order to assess whether the ASO-induced correct splicing leads to the generation of full-length and mature CFTR proteins, protein extracts were prepared and Western blotting was performed using the M3A7 anti‐CFTR antibody. This antibody recognizes the end of the NBD2 domain, enabling the detection of CFTR proteins translated only from correctly spliced transcripts. As can be seen in Figure 1D, treatment with ASO84-1 or ASO84-4 enhanced the production of a full-length, fully glycosylated mature CFTR protein (~175kDa, band C). Altogether, these results clearly show that masking the junctions of the cryptic exon by ASO84-1 and ASO-4 can effectively augment correct CFTR splicing in cells over-expressing the 3849+10kb C-to-T mutation, as reflected both at the mRNA level and protein processing.

### ASO treatment modulates splicing and restores CFTR function in HNEs derived from patients carrying the 3849+10kb C-to-T mutation

For studying the ASO effect on the endogenous CFTR function, we used primary HNEs derived from a CF patient heterozygous for the 3849+10kb C-to-T and the W1282X mutations. ASOs were delivered by free uptake. Short-circuit-current (Isc) experiments, which allow quantification of CFTR-mediated Cl^−^ secretion following forskolin/3-isobutyl-1-methylxanthine (IBMX) activation, showed a significant increase in CFTR channel activity following treatment with 200nM ASO84-1 or ASO84-4 (Figure 2A, 2B). As the W1282X allele does not generate any CFTR-dependent CI^−^ transport (32), the observed current following control ASO treatment reflects low residual CFTR function. Importantly, ASO84-1 and ASO84-4 treatment restored the CFTR channel activity to ~25% of the WT level (32).

**Figure 2.**
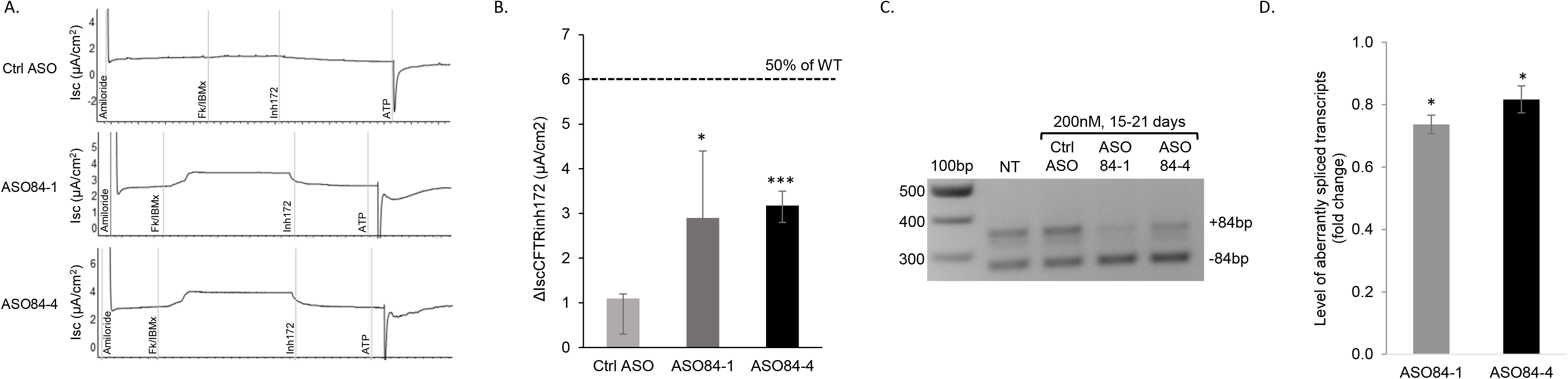
ASO mediated correction of the CFTR splicing pattern and function in primary HNEs carrying the 3849+10kb C-to-T and the W1282X mutations. **A.** Representative traces of Isc measurement by Ussing chamber, in well-differentiated primary HNEs derived from a heterozygote CF patient carrying the 3849+10kb C-to-T/W1282X genotype. The HNEs were cultured in an air liquid interface and treated with 200nM ASO for 17-19 days by free uptake. **B.** Short circuit current values, in response to CFTR specific inhibition (ΔISC CFTR inh172), were measured by Ussing chamber, as described in A. The ASO effect is presented as the median (with min-max range) of the absolute ΔISC CFTR inh172 (μA/cm2) values. Isc was measured from 3 filters treated with a control ASO and 4 filters for each of the ASO84-1 and ASO-84-4 conditions, in 3 independent experiments. Dashed line represents the level of 50% of WT, as described in Pranke et al., (32). Statistical analysis was performed using unpaired t test (two-tail). *p<0.05, ***p<0.001. **C.** RNA was extracted from the HNE filters following the functional analysis and RT-PCR using primers located in exons 22 and 23 was performed to amplify the CFTR transcripts with (+84 bp) and without (−84 bp) the cryptic exon. A representative RT-PCR example is presented (n=3). **D.** Quantitative analysis of the level of aberrantly spliced transcripts was performed by RT-qPCR. The values shown are the average fold change (mean±SEM) of 3 independent experiments relative to cells treated with a control ASO. Values were normalized against transcripts of GUSb gene. Statistical analysis was performed using Student’s t test (one-tail, paired). *p<0.05. NT-non treated.

We next extracted RNA from the HNE filters used for CFTR functional measurements, and analyzed the splicing pattern by RT-PCR and RT-qPCR analyses. ASO treatment modulated the splicing pattern and led to an increase in the relative level of correctly spliced transcripts (Figure 2C). The quantitative analysis by RT-qPCR showed a 30% and 20% reduction in the level of aberrantly spliced CFTR transcripts following treatment with ASO84-1 and ASO84-4, respectively (Figure 2D). These results indicate that ASO splicing modulation is the molecular basis underlying the restoration of the CFTR channel activity in the patient-derived HNEs. Since well-differentiated primary HNEs serve as an *in-vitro* preclinical cellular model able to predict patient-specific respiratory improvement (32, 33), these results demonstrate the clinical potential of ASO based therapy for CF patients carrying the 3849+10kb C-to-T mutation

### ASO-based drug development for CF patients carrying the 3849+10Kb C-to-T mutation

In order to proceed toward clinical development of an ASO-based therapy for patients carrying the 3849+10kb C-to-T allele, we performed the following experiments:

#### I. ASO design and screening for the identification of lead ASO candidates

As the first step we have developed an in-house algorithm which predicts ASO hybridization potency, taking into consideration target coverage, potential immunogenicity and potential off-target effects. Using this algorithm, we designed 26 consecutive ASOs, which align along the entire 84bp cryptic exon and 50bp flanking intronic sequences from both its sides (Figure 3A). The ASOs were initially synthesized with the 2nd generation 2’-OMe/PS chemical modifications and were 18-22nt long. For the establishment of a screening system with a consistent CFTR expression, we introduced a Flp-In acceptor site into Fischer Rat Thyroid (FRT) epithelial cells. The CFTR-3849-mut or CFTR-3849-WT plasmids were integrated into the FRT cell genome via the Flp-In target site (see Materials and Methods for details). For analysis of the ASO effect, FRT-3849-mut cells were transfected with 10nM of each ASO or with a control ASO. Following transfection, RNA was extracted and the CFTR splicing pattern was analyzed by RT-PCR. The results showed that all 26 ASOs tested significantly modified the splicing pattern, resulting in reduced relative levels of aberrantly spliced transcripts accompanied with increased levels of correctly spliced transcripts (Figure 3B). Quantitative analysis by RT-qPCR showed that all ASOs reduced the level of aberrantly spliced CFTR transcripts relative to control ASO, with a fold change ranging from 0.70 for SPL84-26 to 0.20 for SPL84-23 (Figure 3C). Five ASOs (SPL84-2, SPL84-17, SPL84-22, SPL84-23 and SPL84-25) with the highest and most reproducible effect in RT-PCR and in the RT-qPCR were chosen for further analyses (lead ASO candidates).

**Figure 3.**
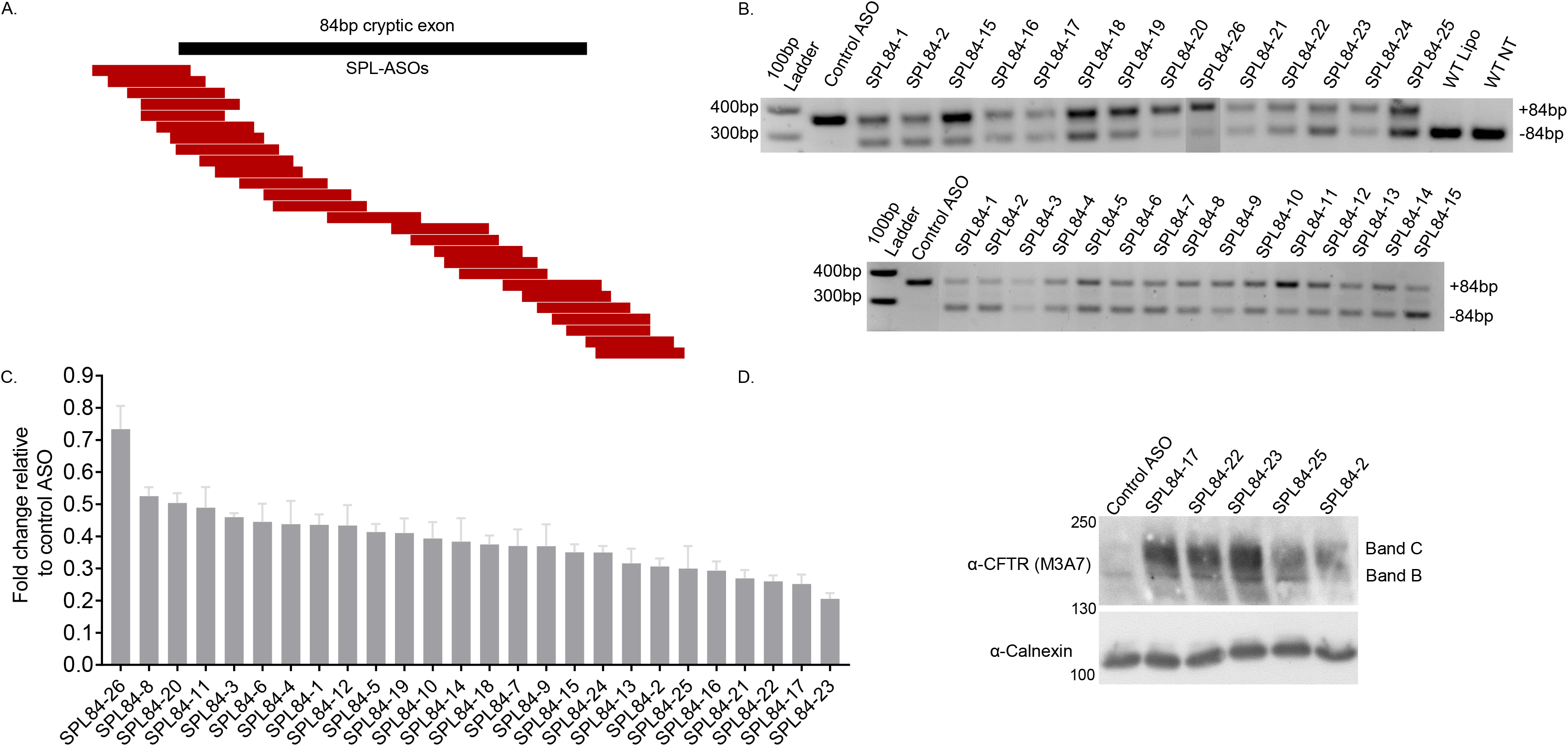
ASO mediated restoration of the CFTR splicing and full-length mature CFTR proteins in FRT cells over-expressing the 3849+10kb C-to-T mutation. **A.** Diagram of the 26 designed ASOs, mapped to their positions relative to the 84bp cryptic exon (black bar) and flanking intronic sequences. **B.** FRT-3849-mut cells were transfected with 10nM of each ASO (named SPL84-X) targeted to consecutive positions along the entire cryptic exon or a control ASO. Twenty-four hours following transfection, RNA was extracted and RT-PCR was performed using primers mapped to exons 22 and 23. Representative examples of RT-PCR from two independent experiments showing the aberrantly (+84bp) and correctly (−84bp) spliced CFTR transcripts. **C.** The level of aberrantly spliced CFTR transcripts was also measured by RT-qPCR performed on RNA extracted following the ASO screen. The values shown are the average fold change (mean±SEM) from 2-8 independent experiments relative to cells treated with a control ASO. Values were normalized against transcripts of the HPRT gene. Paired T test statistical analysis was performed between the delta Ct of each ASO and the delta Ct of the control ASO. The level of aberrantly spliced transcripts following treatment with each ASO was significantly different from the level following control ASO treatment (p<0.05). **D.** FRT-3849-mut cells were transfected with 10nM of each of the five lead ASO candidates or control ASO every 24 hours for 2 days. Protein extracts were prepared and analyzed by immunoblotting with anti CFTR (M3A7) and anti‐calnexin antibodies.

We next analyzed the half-maximal effective concentration (EC50) of the lead ASO candidates by creating a dose-response correlation by transfection of FRT-3849-mut cells with six concentrations of each ASO, ranging between 0.1nM-100nM. Following RNA extraction and CFTR splicing analysis, the EC50 and maximal effect (efficacy) were calculated for each ASO. As presented in Supplementary Figure 2, the ASO treatments resulted in a dose-dependent decrease in the level of aberrantly spliced CFTR transcripts, with very low EC50 values (<1nM) for all five lead ASO candidates, highlighting their high potency and efficacy.

#### II. The effect of the lead ASO candidates on CFTR protein production and maturation in FRT and HEK293 cells

To analyze the effect of the lead ASO candidates on the production of mature CFTR protein, we analyzed their effect in FRT-3849-mut cells. As can be seen in Figure 3D, almost undetectable levels of full-length mature protein can be seen after treatment with a control ASO. Following treatment with the lead ASO candidates, the full-length mature CFTR protein (band C) was clearly observed (Figure 3D). The level of full-length mature CFTR proteins differed among the various ASOs, with the best effect achieved with SPL84-17, SPL84-22 and SPL84-23. In order to verify this result, the effect of the lead ASO candidates on CFTR protein production was analyzed in an additional cellular system, the HEK293-3849-mut cells, which presented similar results (Supplementary Figure 3D). Altogether, the selected ASOs modulate the CFTR splicing pattern and lead to the formation of a full-length and mature CFTR protein.

#### III. The effect of the lead ASO candidates on CFTR function in primary nasal and bronchial epithelial cells from CF patients carrying the 3849+10Kb C-to-T mutation

Final lead ASO selection was performed by analyzing the effect of the lead ASO candidates (SPL84-2, SPL84-17, SPL84-22, SPL84-23 and SPL84-25) on CFTR activity in primary HNEs, using Isc experiments. The ASOs were introduced into the cells by free uptake. We analyzed HNEs derived from five CF patients, heterozygous for the 3849+10Kb C-to-T mutation and various minimal function mutations (F508del, W1282X or 405+1G-to-A) and one patient homozygous for the 3849+10Kb C-to-T mutation. The analysis of HNEs derived from the homozygous patient (Figure 4A and 4B) showed a low baseline CFTR activity. Treatment with each of the lead ASO candidates led to a complete rescue of CFTR function, showing the significant high potency of all lead ASOs (Figure 4B).

**Figure 4.**
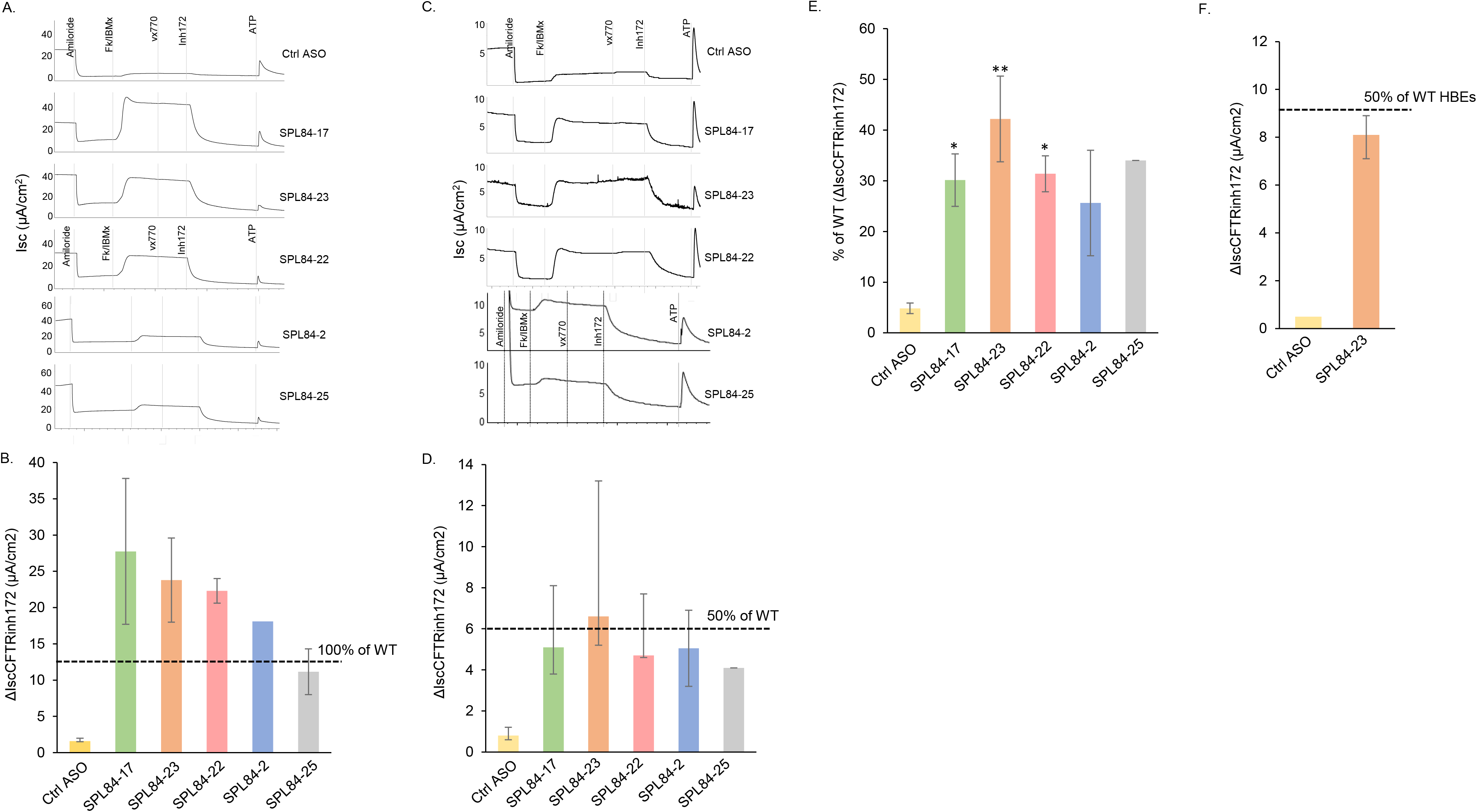
Lead ASO candidates rescue CFTR function in well-differentiated primary HNEs and HBEs derived from patients heterozygous or homozygous for the 3849+10kb C-to-T mutation. **A** and **C.** Representative traces of electrophysiological responses, measured by Ussing chamber, in HNEs derived from a homozygote (**A**) or a heterozygote (3849+10kb C-to T/F508del) (**C**) patient, following treatment with the indicated ASO for 15-21 days. **B** and **D.** The effect of each ASO on HNEs derived from the same patients as in A and C is presented as the median (with min-max range) of the absolute ΔIsc CFTR inh172 values. **B.** The measurements are from filters treated with SPL84-17, SPL84-23, SPL84-22, SPL84-25 (n=2 for each ASO), SPL84-2 (n=1), control ASO (n=3). **D.** The measurements are from filters treated with the control ASO (n=5), SPL84-22, SPL84-23, SPL84-17 (n=4 for each ASO), SPL84-2 (n=2), SPL84-25 (n=1). **E.** Mean CFTR activation values are presented as % of WT CFTR activity [Pranke et al., (32)] following ASO treatment in HNEs from 5 heterozygote patients. Mean (±SEM) response to each ASO was calculated from the median values of all patients. The values were calculated from filters treated with the control ASO (n=21, derived from 5 patients), SPL84-17 or SPL84-22 (n=9 for each ASO) derived from 3 patients, SPL84-23 (n=24, derived from 5 patients), SPL84-2 (n=5, derived from 2 patients) and SPL84-25 (n=1, derived from one patient). Statistical analysis was performed using paired t test (two-tail). *p<0.05, **p<0.01. **F.** The effect of SPL84-23 on HBEs derived from a heterozygous patient (3849+10kb C-to-T/F508del) is presented as the median (with min-max range) of the absolute ΔIsc CFTR inh172 values, calculated from 3 filters. The horizontal dashed line indicates WT levels in HNEs (B, D) or HBEs (F) according to Pranke et al., (32). In all experiments 200 nM ASO was used.

We next analysed the ASO effect in HNEs from patients heterozygous for the 3849+10kb C-to-T mutation. An example of the ASO effect on one of the patients, with the 3849+10kb C-to-T/F508del genotype, is presented in Figure 4C and 4D. As can be seen, the baseline CFTR activity level in this patient was very low but treatment with each of the lead ASOs led to a significant CFTR activation, reaching 34-55% of WT activity (Figure 4D). These functional results highlight the major effect of SPL84-23 which was able to restore the CFTR activity to 55% of WT, the level found in healthy individuals carrying a CFTR mutation, reflecting full restoration of CFTR activity from the 3849+10kb C-to-T allele (32). Analysis of the average ASO effect in HNEs derived from five heterozygous patients showed a significant CFTR activation of each lead ASO candidate ranging from 26% of WT level for SPL84-2 to 43% for SPL84-23 (Figure 4E). As a control for the specificity of SPL84-23 to the 3849 allele, we analyzed the ASO effect on the CFTR function in HNEs derived from a patient homozygous for the F508del mutation. As can be seen in Supplementary Figure 4 the cells had no CFTR activity following forskolin treatment, with and without the addition of the ASO, indicating that the splicing modulation and CFTR activation by SPL84-23 is specific to the 3849 allele. Altogether, based on the results from the screening systems and the human primary nasal cells, ASO SPL84-23 was selected as the lead ASO for further development and clinical assessment. Treatment of primary HBEs (derived from explanted lungs from a CF patient carrying the 3849+10kb C-to-T/F508del mutations) with the lead ASO SPL84-23 led to a marked rescue of CFTR function, reaching a CFTR activity level of 43% of WT (Figure 4F). These results further confirm the high therapeutic potential of SPL84-23 to rescue CFTR function in patients carrying the 3849+10kb C-to-T allele. Well differentiated cultured HBEs exhibit many of the morphological and functional characteristics believed to be associated with CF airway disease *in vivo* [reviewed in (34, 35)] and serve as a gold standard for drug development in CF.

It is worth noting that treatment with VX-770, which is approved for patients carrying the 3849+10kb C-to-T mutation (36), had a minimal effect on CFTR activity in HNEs from a patient carrying the 3849+10kbC-to-T/F508del genotype (Supplementary Figure 5). In addition, treatment with VX-770 combined with a control ASO in HNEs from another patient carrying the 3849+10kb C-to-T/F508del genotype also showed a minimal effect (Figure 4A and 4C). Treatment with each of the lead ASO candidates had no apparent additional effect with VX-770 versus ASO alone (Figure 4A and 4C). These results highlight the need for an efficient therapy for patients carrying the 3849+10kb C-to-T mutation and further demonstrate the therapeutic potential of ASOs for these patients.

#### IV. The effect of the lead ASO SPL84-23 on the *splicing pattern* in primary HNE cells from CF patients carrying the 3849+10Kb C-to-T mutation

We further analyzed the effect of SPL84-23 on the splicing pattern in the well differentiated heterozygous and homozygous HNEs that were analyzed in the functional assay presented in Figure 4. As can be seen in Figure 5A, in HNEs derived from the homozygous patient the majority of the transcripts are aberrantly spliced. SPL84-23 treatment significantly redirected the splicing pattern towards correct splicing. To evaluate the effect of ASO SPL84-23 on the splicing pattern in heterozygous patients, we used HNEs derived from patients carrying the W1282X mutation on the second allele. In these cells, due to NMD, the expression of correctly spliced transcripts from the W1282X allele is markedly diminished. As seen in Figure 5B, in these cells SPL84-23 also significantly redirected the splicing pattern towards correct splicing. Quantitative analysis of RNA derived from filters of 4 heterozygous and the homozygous patients showed a fold change of ~0.2-0.3 in the level of aberrantly spliced transcripts relative to the control ASO, indicating that the marked rescue of CFTR function by SPL84-23 treatment resulted from the major decrease in aberrantly spliced transcripts (Figure 5C). Altogether, these results indicate that the ASO mode of action, underlying the significant CFTR functional correction, is the generation of correctly spliced transcripts.

**Figure 5.**
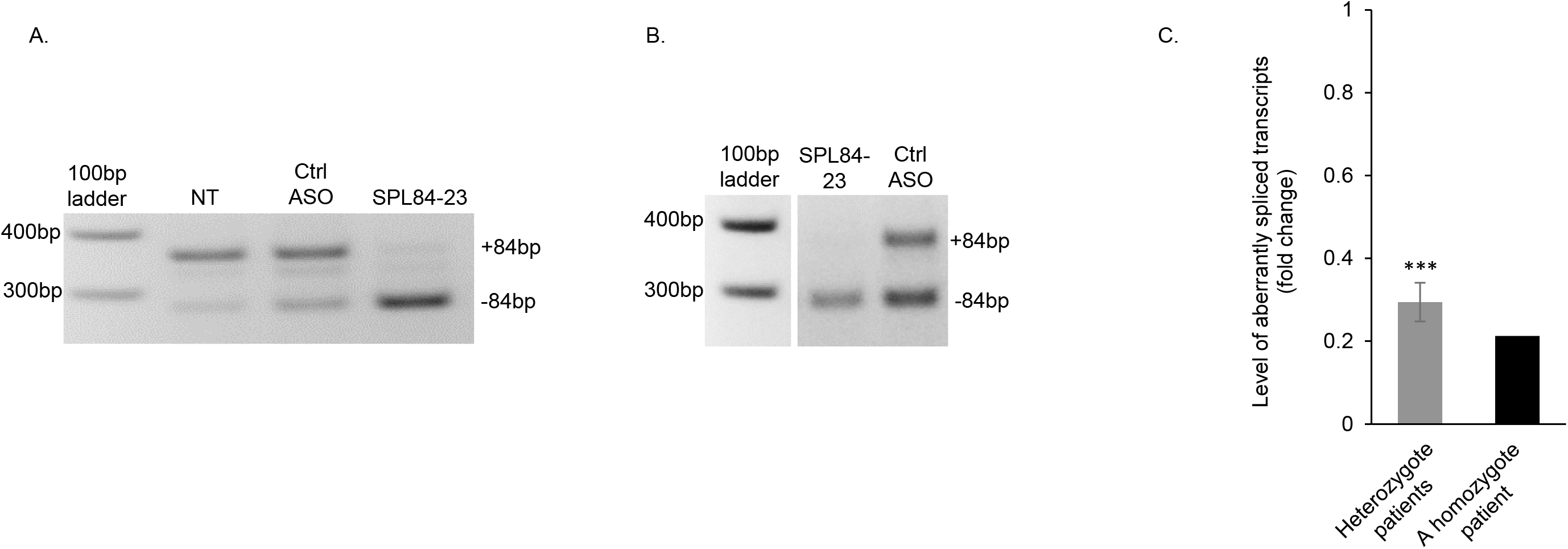
Splicing modulation by the lead ASO SPL84-23 in HNEs derived from homozygous and heterozygous patients. RNA was extracted from HNE cells derived from patients heterozygous (N=4, 7 filters) or homozygous (N=1, one filter) for the 3849+10kb C-to-T mutation, following the functional measurements presented in Figure 4. The effect on the CFTR splicing pattern was analyzed using RT-PCR and RT-qPCR. Representative RT-PCRs examples of the aberrantly and correctly spliced CFTR transcripts following treatment with SPL84-23 in HNE cells derived from the homozygous patient (**A**) or the heterozygous patient (3849+10kb C-to-T/W1282X) (**B**) are shown. RT-PCR using primers located in exons 22 and 23 was performed to amplify the aberrantly spliced CFTR transcripts (+84 bp) and the correctly spliced (−84bp). **C.** The effect of SPL84-23 on the level of aberrantly spliced CFTR transcripts as measured by RT-qPCR. The values shown are the fold change relative to cells treated with a control ASO. Values were normalized against transcripts of GUSb gene. Statistical analysis was performed using paired t test (one-tail). ***p<0.001. NT-non treated.

### V. Optimization of SPL84-23 potency by chemical modifications

The experiments described above were performed using ASOs containing 2’-OMe/PS chemical modifications. For further optimizing the SPL84-23 efficiency we evaluated the 2’-Methoxy Ethyl (2’-MOE) sugar modification on the background of a full phosphorothioate backbone (2’-MOE/PS). This 2’-MOE chemical modification confers increased nuclease resistance and higher affinity to the target RNA (30). In order to compare the effect of 2’-OMe/PS to 2’-MOE/PS chemical modifications, CFTR functional analysis was performed on primary HNEs derived from a patient homozygous for the 3849+10Kb C-to-T mutation, following exposure by free uptake to SPL84-23 with the two different modifications. For this the cells were treated with two ASO concentrations, 50nM and 200nM, of each chemistry. Treatment with SPL84-23 carrying the 2’-OMe chemistry presented a dose response with full functional restoration following 200nM treatment (Figure 6A, green bars and Supplementary Figure 6A). Strikingly, full CFTR functional restoration was achieved following treatment with SPL84-23 carrying the 2’-MOE modification already at 50nM, which showed a further increase following treatment with 200nM (Figure 6A blue bars and Supplementary Figure 6B). These results clearly show the benefit of the 2’-MOE modification in ASO-mediated restoration of CFTR function. Analyzing the effect of the different chemically modified ASOs on the splicing pattern supported the functional results. The effect of SPL84-23 carrying the 2’-MOE modification on the splicing pattern was more efficient than that of the 2’-OMe modification, as demonstrated by a significant reduction in the level of aberrantly spliced transcripts already at 50nM, that was further enhanced by 200nM (Figures 6B and 6C). Similar results showing a clear benefit of the 2’-MOE chemistry over the 2’-OMe chemistry in correcting the splicing pattern were observed also in the FRT-3849-mut cells following transfection of 10nM SPL84-23 carrying the different chemical modifications (Supplementary Figure 7).

**Figure 6.**
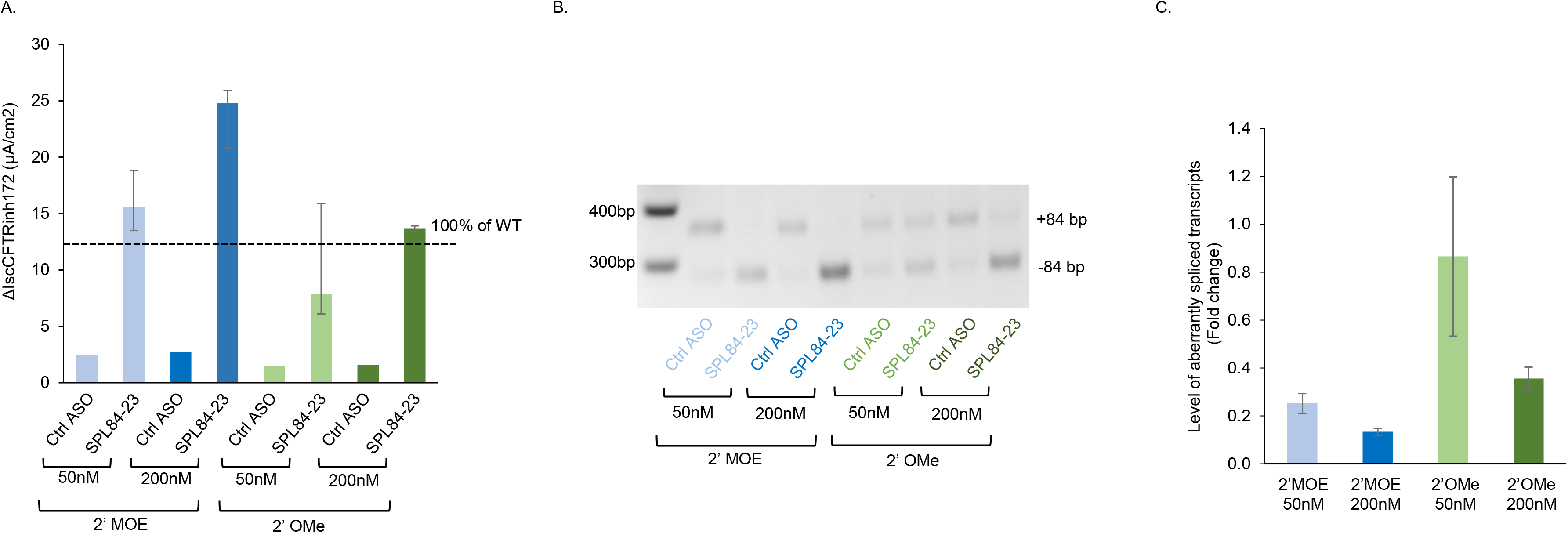
The 2’-MOE chemical modification increases ASO SPL84-23 efficiency in HNEs derived from a CF patient homozygous for the 3849+10kb C-to-T mutation. Well-differentiated primary HNE cells derived from a patient homozygous for the 3849+10kb C-to-T mutation were treated with 50nM or 200nM of ASO SPL84-23, synthesized with two different chemical modifications, 2’MOE and 2’OMe. **A.** The ASO effect is presented as the median (with min-max range) values of Isc, in response to CFTR inh172 measured by Ussing chamber from 2-3 filters treated with each of the SPL84-23 concentrations and from one filter treated with each of the control ASO concentrations. The horizontal dashed line indicates the average level of ΔIsc CFTR inh172 in HNE cultures from healthy WT individuals, according to Pranke et al., 2017 (32). **B** and **C.** RNA was extracted from the HNE filters following the functional analyses. The aberrantly and correctly spliced CFTR transcripts were analyzed by RT-PCR using primers located in exon 22 and 23 (**B**) and quantitative analysis of the level of aberrantly spliced transcripts was performed by RT-qPCR (**C**). The shown values are the average fold change (mean±SEM) following treatment with the indicated chemically modified ASO (2-3 filters from each treatment) relative to cells treated with a control ASO (one filter from each treatment). Values were normalized against transcripts level of GUSb.

Altogether, SPL84-23 carrying the 2’-MOE modification presented superiority over the 2’-OMe modification in correcting CFTR splicing and function. Thus, SPL84-23 with the 2’-MOE modification was selected as the lead ASO for further drug development.

## Discussion

In the present study, we have described a developmental path which led to the identification of a lead ASO drug candidate able to correct the aberrant splicing and restore normal full-length functional CFTR protein in primary, well-differentiated HNEs and HBEs from patients carrying the 3849+10kb C-to-T mutation. Our results clearly show that the lead ASO drug candidate is highly effective and potent, displaying a significant and consistent splicing modulation effect which leads to restoration of CFTR function, reaching levels expected to confer a significant clinical benefit and improved quality of life for patients carrying this splicing mutation [reviewed in (37)].

The ability of splicing modulation to rescue CFTR protein processing and function in cells carrying the 3849+10kb C-to-T mutation was previously addressed by us and others (21, 24, 26). We have previously shown that over-expression of splicing factors in nasal epithelial cells carrying the 3849+10kb C-to-T mutation increased the level of correctly spliced transcripts and restored CFTR channel function (21). A recent study further supported these findings, showing that transfection of a 25-mer PMO (targeting the donor 5’ splice site generated by the 3849+10kb C-to-T mutation) was able to reduce aberrant splicing and to improve CFTR channel activity in HBEs derived from CF patients carrying at least one allele of the 3849+10kb C-to-T mutation (26). In line with this important recent data, our present work, performed both in cells over-expressing the 3849+10kb C-to-T mutated CFTR allele and in respiratory primary cells derived from CF patients homozygous for the mutation or heterozygous with various second alleles, establishes that splicing modulation is a powerful tool for restoration of CFTR in cells carrying a splicing abnormality. In the current work, we describe development of a highly potent ASO-based drug candidate, a 2’-MOE/PS modified 20nt ASO, that was efficiently delivered by free uptake into well differentiated HNEs and was able to completely restore correct splicing and function at low nanomolar concentrations. The lead ASO candidate rescued CFTR expression and function to WT levels in HNEs from a homozygous patient and reached an average of 43% of WT levels in HNEs and HBEs from six heterozygous patients, 5 of which achieving levels ≥37% of WT. Correlations between CFTR functional modulation in patient-derived HNE cells with clinical efficacy showed that reaching CFTR-dependent Cl^−^ secretion levels >10% of WT function corresponds to a significant improvement in the respiratory activity measured as change in ppFEV1 (32). Indeed, exceeding 10% of WT CFTR activity is associated with pancreatic sufficiency and less elevated sweat chloride levels [reviewed in (38)]. It is worth noting that, to the best our knowledge, our ASO-based therapy is the first approach leading to the generation of normal full-length functional CFTR proteins. Our results therefore provide a solid basis for advancement of our lead ASO, expected to confer significant clinical benefit.

It is important to note that ASOs with 2’-MOE chemistry are already in clinical use. The FDA-approved SPINRAZA^®^, a splice-switching ASO indicated for SMA, composed of the 2’-MOE modification, that demonstrates high efficacy in patients with early and later-onset disease (39). The favorable benefit-to-risk profile of this and other 2’-MOE-based drugs (30), prompted us to evaluate the efficacy of our lead ASO with 2’-MOE chemical modification. As presented in the current study, enhanced ASO efficiency was achieved by the 2’-MOE chemical modification, which allowed the use of significantly reduced concentrations for complete CFTR restoration (Figure 6).

Based on our results, we aim to develop an inhaled ASO drug, for increased lung exposure with minimal systemic exposure. Previous studies showed that inhaled ASOs are widely distributed and active in mice and non-human primates lungs, reaching even the most distal (alveolar) regions, with minimal systemic exposure and good tolerability (40–43). Importantly, in a recent study, Crosby et al. demonstrated that ASOs dissolved in saline traverse CF-like mucus and distribute throughout mouse lung (44). Moreover, the ASOs were found to be effective and lead to a reduction of the target mRNA. These studies support the notion that inhaled ASO-based drugs can be efficiently delivered to the lungs of CF patients.

The 3849+10kb C-to-T mutation is one of the ten most common CFTR mutations, carried by >1400 individuals worldwide. The CFTR modulators Kalydeco® (Ivacaftor/VX-770) and Symdeco® (Tezacaftor/Ivacaftor) were approved recently in some countries for CF patients carrying the 3849+10kb C-to-T splicing mutation. While both ivacaftor and tezacaftor/ivacaftor are clinically approved for the treatment of people with CF caused by the 3849+10kb C-to-T mutation, with or without the F508del mutation, clinical trials demonstrate only modest response compared to placebo. Changes in spirometry (i.e. FEV_1_) ranged from ~2.6-5.8% (12, 13, 36, 45), far short of the benchmark set by highly effective CFTR modulator therapy (46, 47), leaving an unmet need for a more effective therapy. Indeed, as presented in the current paper, *in vitro* treatment of HNEs heterozygous for the 3849+10kb C-to-T mutation with VX-770 alone, resulted in a minimal CFTR response (Supplementary Figure 5), noting the high degree of cAMP stimulation by forskolin that preceded VX-770 administration may have abrogated the modest effect, as reported previously (48). Furthermore, the addition of VX-770 to our highly effective lead ASO candidates showed no additional effect to that of the ASO alone (Figure 4A and 4C). These results support recent data showing small and inconsistent improvement in chloride transport following VX-770 treatment of HNEs from CF patients heterozygous and homozygous for the 3849+10kb C-to-T mutation (26). Given the limited clinical benefit of the currently available CFTR modulators for patients carrying the 3849+10kb C-to-T mutation, the ASO strategy presented here, which leads to the generation of normal CFTR protein, has the potential to more affirmatively restore CFTR function and provide a potential significant clinical benefit to patients with the 3849+10kb C-to-T mutation. As other CFTR splice variants, e.g., 2789+5 G-to-A, 3272-26 A-to-G and 1811+1.6 kb A-to-G, have a mechanistically similar splicing defect, the concept is likely to apply to additional CFTR alleles with potential benefit for these patients.

In summary, the results presented here highlight the therapeutic potential and clinical benefit of ASO-based splicing modulation for genetic diseases caused by splicing mutations and pave the way towards clinical assessment of our lead ASO.

## Materials and Methods

### CFTR plasmids

CFTR expression plasmids (CFTR-3849-mut and CFTR-3849-WT) were constructed with the full length CFTR cDNA into which sequences from intron 22 including the 84 bp cryptic exon, the 3849+10kb C-to-T mutation or the WT sequence and relevant flanking intronic sequences were inserted (49). CFTR-3849-WT and CFTR-3849-mut cDNA were cloned into pcDNA5/FRT, an expression vector designed for use with the Flp-In System (Invitrogen) (Supplementary Figure 1).

### Cell lines and cell transfections

HEK293 were grown in DMEM supplemented with 10% fetal calf serum. For ASO splicing modulation, HEK293 cells were transiently transfected with the CFTR-3849-mut plasmid for 48 hours (HEK293-3849-mut), followed by a second transfection of 100nM ASO 24 hours after the initial transfection with the CFTR plasmid. RNA and protein were extracted 24 hours after ASO transfection.

For the establishment of a screening system with a consistent CFTR expression, FRT cell models were generated. FRT cells were stably transduced with the CFTR-3849-mut and matched CFTR-3849-WT control cDNA using the Flp-In™ system per the manufacturer’s protocol (Invitrogen). The cells were grown and maintained in Nutrient Mixture F-12 Ham (Sigma-Aldrich) medium supplemented with 2.68 g/L sodium bicarbonate and 5-10% FBS as previously described (50). For the ASO screen, FRT-3849-mut cells were transfected with 10nM of each of the ASOs and RNA was extracted after 24 hours. For the EC50 experiment, in order to ensure uniform transfection conditions across all different ASO concentrations, we maintained a constant ASO concentration of 100nM across all points of the dose-response curve by adding a complement amount of control ASO to the tested ASO. For protein evaluation, FRT-3849-mut cells were transfected every 24 hours with 10nM of each ASO and total protein was extracted following 72 hours from the first transfection. Lipofectamin 2000 Transfection Reagent (Thermo Fischer scientific) was used for transfecting the various ASOs. TransIT®-LT1 Transfection Reagent (Mirus) was used for transfecting the plasmids

### RNA analysis of splicing pattern

Total RNA from the cell lines was extracted using the RNeasy mini Kit (Qiagen). Total RNA from HNEs or HBEs was extracted using the QIA shredder and RNeasy micro Kit (Qiagen). Complementary DNA synthesis was performed using the High Capacity cDNA kit (Applied Biosystems). For the evaluation of the cryptic exon skipping, we employed two methods: (1) RT-PCR to amplify the correctly and aberrantly spliced transcripts using primers aligned to exon 22 and 23, using Platinum™ SuperFi™ Green PCR Master Mix (Invitrogen). PCR Primers: F-5’-ATAGCTTGATGCGATCTGTGA-3’ (exon 22) and R-5’-ATCCAGTTCTTCCCAAGAGGC-3’ (exon 23). (2) RT-qPCR for quantitative detection of aberrantly spliced CFTR transcripts containing the 84bp cryptic exon using TaqMan master Mix (Applied Biosystems). The expression level was normalized to the transcript levels of HPRT (for FRT cells) or GUSb (for human cells). For statistical analysis of the ASO effect, paired t-test was performed on the delta Ct values of each ASO relative the delta Ct values of the control ASO.

### Primary nasal and bronchial epithelial cell sampling and culture

Bilateral nasal brushing was performed to obtain cells from the lower turbinate in the middle third from CF patients, heterozygous or homozygous for the 3849+10kb C-to-T mutation. The brushes were immediately immersed in PBS. HBE cells were derived from lung explants of a CF patient heterozygous for the 3849+10kb C-to-T mutation. Lung tissue specimens obtained from lungs that were removed during lung transplantation, were transported to the laboratory in a container filled with PBS or culture medium, on wet ice. For increasing the number of cells, freshly isolated HNE and HBE cells were expanded on flasks using conditionally reprogramming media (32). The initial expansion phase was maintained for 2-3 passages. After expansion, cells were seeded on porous filters (Transwell Permeable Supports, 6.5 mm Inserts, 24 well plate, 0.4 μm Polyester Membrane, ref.: 3470, Corning Incorporated) for differentiation in a liquid-liquid interface for 2 - 5 days. After that period the apical medium was removed and air liquid interface culture was initiated and maintained for further 15-23 days, a period which was previously found to allow polarization and differentiation of respiratory epithelium (32). The ASOs were added to the HNE or HBE cells by a free uptake during medium exchange, every other day.

### Ussing chamber measurements on HNEs and HBEs

The Isc was measured under voltage clamp conditions with an EVC4000 precision V/I Clamp (World Precision Instruments). Culture inserts with differentiated HNE or HBE cells were mounted in Ussing chambers (Physiologic Instrument, San Diego, CA). For all measurements, Cl^−^ concentration gradient across the epithelium was applied by differential composition of basal and apical Ringer solutions. Inhibitors and activators were added after stabilization of baseline Isc: sodium (Na^+^)-channel blocker Amiloride (100 μM) to inhibit apical epithelial Na+ channel (ENaC); cAMP agonists forskolin (10 μM) and IBMX (100 μM) to activate the transepithelial cAMP-dependent current (including Cl^−^ transport through CFTR channels); VX-770 (10 μM) or genestein (10 μM) to potentiate CFTR activity, CFTR inhibitor CFTRinh172 (5 μM) to specifically inhibit CFTR and ATP (100 μM) to challenge the purinergic calcium-dependent Cl^−^ secretion. CFTR specific activity was quantified according to the current change following the addition of the CFTR inhibitor Inh-172, which enables to determination of the relative current contribution of the CFTR channels versus other anion transport pathways (51). These parameters served as an index of CFTR function. The data from each patient is presented in median values. At least 2 filters were analyzed per patient to assess reliable results.

### Western Blot

Protein extracts were prepared using RIPA buffer. 6% Polyacrylamide gels were used for protein separation. The gel was transferred to a nitrocellulose membrane and antibody hybridization and chemiluminescence were performed according to standard procedures. The primary antibodies used in this analysis were mouse anti CFTR M3A7 (Millipore) and rabbit anti Calnexin (Sigma). HRP-conjugated anti-rabbit and anti-mouse secondary antibodies were obtained from Jackson Immunoresearch Laboratories.

### Statistics

Statistical analyses were performed with GraphPad Prism version 9.0 or Excel software. Differences between groups were determined by t-test. For all statistical tests, P values less than 0.05 were considered significant.

### Study approval

CF patients were recruited for HNE sampling (Clinical Trials: NCT02965326). The exclusion criteria were smoking, local nasal treatment and rhinitis at the time of sampling. The study was approved by the Ile de France 2 Ethics Committee, and written, informed consent was obtained from each adult and parent (AFSSAPS (ANSM) B1005423-40, n° Eudract 2010-A00392-37; CPP IDF2: 2010-05-03-3). HBE cells were derived from lung explants after written informed consent. All experiments were performed in accordance with the guidelines and regulations described by the Declaration of Helsinki in France and Israel.

## Author contributions

YSO and MI-TS are co-first authors, with YSO listed first because of her extensive contribution to the project conception and design, experiments, data analyses and manuscript writing; MI-TS initiated the project, contributed to its design, data analyses and led the manuscript writing. OBA contributed to project design, ASO design, conducted experiments and data analyses; AG designed and conducted functional experiments in patient cells; AG and OA are co-second authors, with AG first because she contributed extensively to the functional experiments; AH conducted functional experiments in patient cells; VM together with YL and JH constructed the plasmids and established the ASO screening system; EOG contributed to the project design, ASOs design and data analyses; JR contributed to collecting epithelial cells from patients; EJS contributed to the design of the screening system and obtaining financial support; SDW contributed to ASO design and synthesis; EK contributed to the design of the project, the recruitment of patients and nasal epithelial cells scraping as well as data analyses and discussion; SMR contributed to the conception and design of the ASO screening system, data analysis, obtaining financial support and manuscript writing; ISG contributed to the design and supervision of the functional experiments, recruitment of patients for nasal epithelial cell scraping, data analyses and manuscript writing; BK contributed to the conception and design of the project, data analysis and interpretation, obtained financial support, manuscript writing and final approval of the paper. All authors contributed and edited the manuscript.

## Acknowledgements

We are grateful to all CF patients for their participation in the research. We thank all the CF centers for their support and effort to encourage patients to participate in the research. We thank the entire past and present membership of the Kerem group for technical advice, assistance and enlightening discussions. This research was supported (in part) by grants from the Cystic Fibrosis Foundation to B.K. (KEREM13XX0, KEREM15XX0) and S.M.R. (ROWE19R0), the Israel Ministry of Economy and Industry, the Kamin program to B.K. and the NIH to E.J.S. and S.M.R. (P30DK072482).

