## Supplemental Figures 1-6 for "Antisense oligonucleotide-based drug development for Cystic Fibrosis patients carrying the 3849+10kb C-to-T splicing mutation"

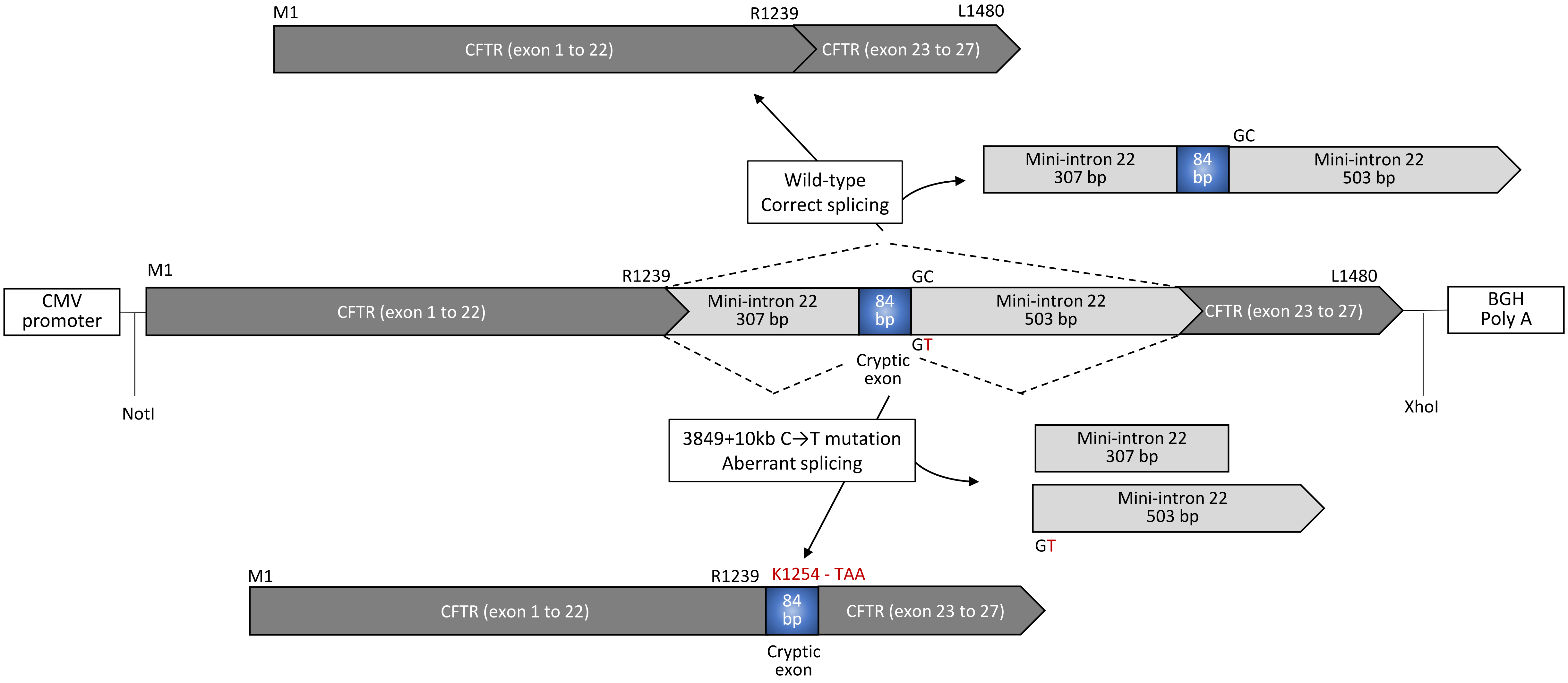


**Supplementary Figure 1. Schematic diagram of 3849+10kb C-to-T CFTR expression plasmid and expected CFTR protein product**. CFTR cDNA was cloned between the NotI and XhoI sites of the pcDNA5/FRT vector. 3849+10kb C-to-T CFTR expression plasmid was constructed by inserting mini-intron 22 behind arginine codon 1239 [Nissim-Rafinia et al., 2000 (49)]. Mini-intron 22 was 894 bp long and consisted of three segments: 307 bp 5' arm, 84 bp cryptic exon, and 507 bp 3' arm bearing the 3849+10kb C-to-T mutation. Control WT plasmid was identical to the mutant plasmid without the C-to-T mutation. The middle diagram represents the expression construct with mini-intron 22. The top diagram represents the correct splicing result in WT CFTR and 1480 amino acid full-length CFTR. The bottom diagram depicts 3849+10kb C-to-T CFTR mRNA with the 84 bp cryptic exon inserted due to aberrant splicing. The cryptic exon inserts 15 amino acids behind R1239 followed by a TAA termination codon.

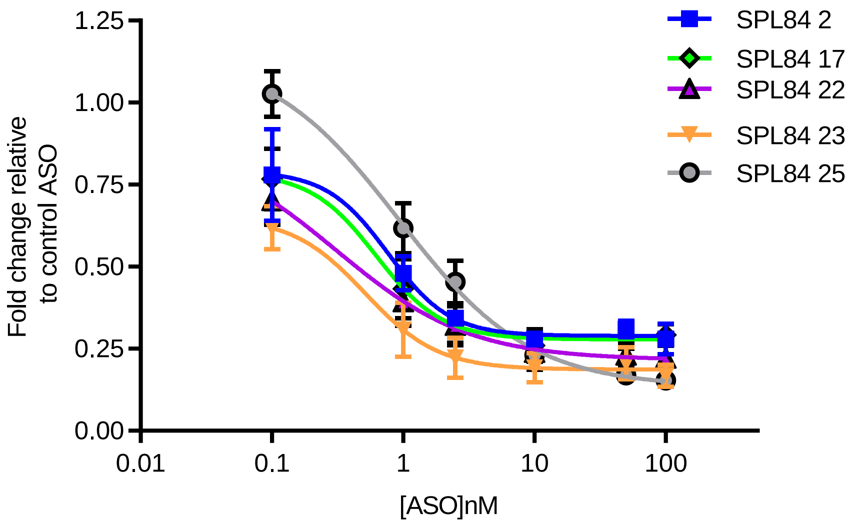


| | SPL84 2 | SPL84 17 | SPL84 22 | SPL84 23 | SPL84 25 |
|---|---|---|---|---|---|
| Maximal effect | 0.29 | 0.28 | 0.22 | 0.19 | 0.13 |
| EC50(nM) | 0.77 | 0.62 | 0.31 | 0.51 | 0.92 |

**Supplementary Figure 2: EC50 and maximal effect of the lead ASO candidates (non-linear regression analysis)** FRT-3849-mut cells were transfected with the indicated concentrations of each ASO. Twenty four hours following transfection, RNA was extracted and quantitative analysis of the level of aberrantly spliced transcripts was performed by RT-qPCR. The values are the average fold change (mean±SEM) from 3-6 independent experiments, relative to cells treated with the control ASO. Values are normalized against transcripts of the HPRT gene.

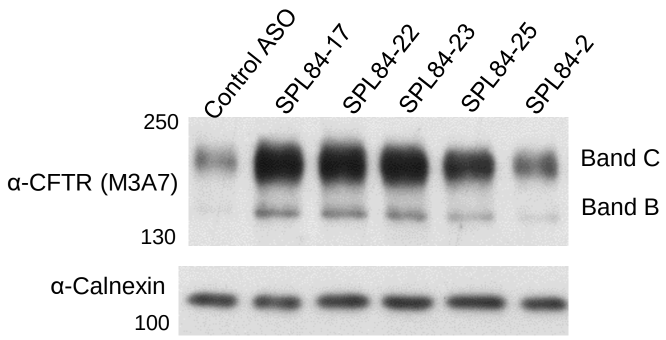


**Supplementary Figure 3: ASO mediated restoration of full-length mature CFTR proteins in HEK293 cells over-expressing the 3849+10kb C-to-T mutation.** HEK293-3849-mut cells were transfected with 100nM of the indicated ASOs for 24 hours. Protein extracts were prepared and analyzed by immunoblotting using anti CFTR (M3A7) and anti-calnexin antibodies.

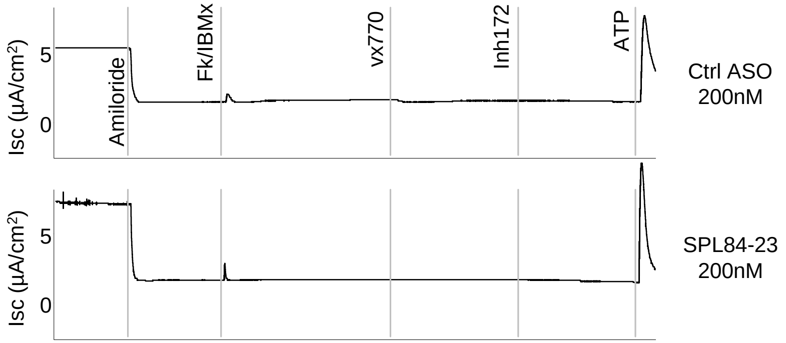


**Supplementary Figure 4: Specificity of SPL84-23 ASO to the 3849+10kb C-to-T mutation.** Representative traces of electrophysiological responses of HNEs derived from a CF patient homozygous for the F508del mutation, measured by Ussing Chamber following control and SPL84-23 ASOs.

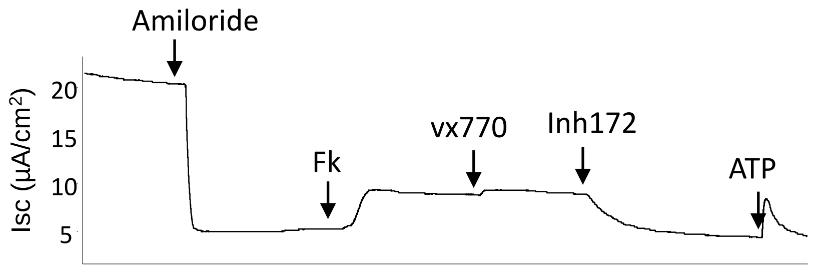


**Supplementary Figure 5: The effect of VX-770 alone on the CFTR function in HNEs derived from a heterozygous patient carrying the 3849+10kb C-to-T and the F508del mutations.** A representative trace of electrophysiological responses of 3849+10kb C-to-T/F508del HNEs measured by Ussing Chamber following forskolin activation, VX-770 treatment and Inh172 inhibition.

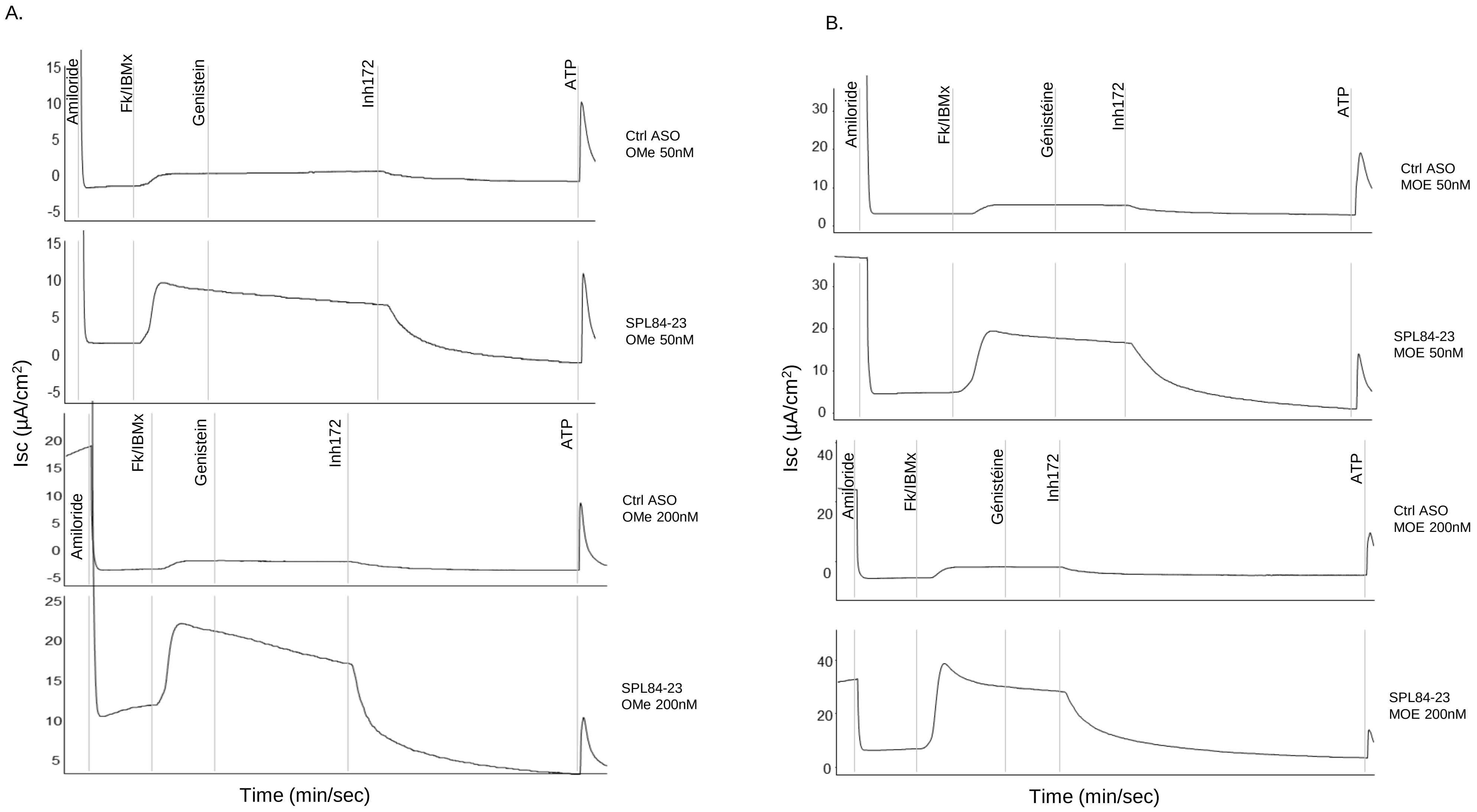


**Supplementary Figure 6: The 2'-MOE chemical modification increases ASO SPL84-23 efficiency in HNEs derived from a CF patient homozygous for the 3849+10kb C-to-T mutation.** Representative traces of electrophysiological responses of 3849+10kb C-to-T/3849+10kb C-to-T HNEs measured by Ussing Chamber following treatment with 50nM or 200 nM ASO SPL84-23 with the 2'-OMe (A) or 2'-MOE (B) chemical modification.

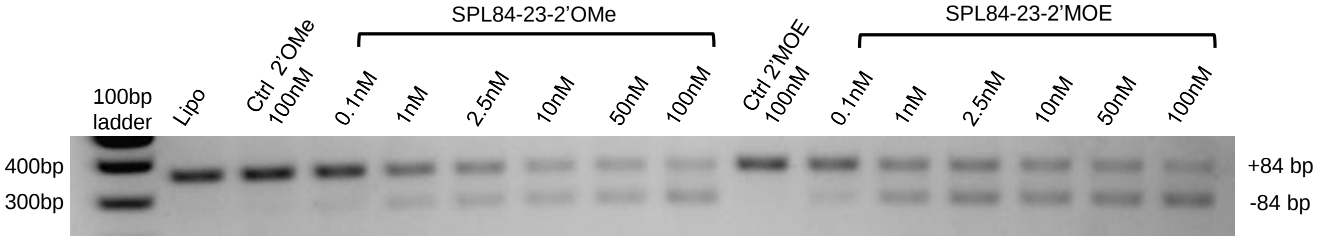


**Supplementary Figure 7: The 2'-MOE chemical modification increases ASO SPL84-23 efficiency in FRT cells over-expressing the 3849+10kb C-to-T mutation.** FRT-3849-mut cells were transfected with the indicated concentrations of SPL84-23, synthesized with two different chemical modifications, 2'-MOE or 2'-OMe. RNA was extracted 24 hours following the transfection and RT-PCR using primers located in exons 22 and 23 was performed to amplify the aberrantly (+84 bp) and correctly spliced (-84bp) CFTR transcripts. A representative example of RT-PCR is shown.
